# Method for Improving Psychophysical Threshold Estimates by Detecting Sustained Inattention

**DOI:** 10.1101/275594

**Authors:** Mike D. Rinderknecht, Raffaele Ranzani, Werner L. Popp, Olivier Lambercy, Roger Gassert

## Abstract

Psychophysical procedures are applied in various fields to assess sensory thresholds. During experiments, sampled psychometric functions are usually assumed to be stationary. However, perception can be altered, for example by loss of attention to the presentation of stimuli, leading to biased data which results in poor threshold estimates. The few existing approaches attempting to identify non-stationarities either detect only whether there was a change in perception, or are not suitable for experiments with a relatively small number of trials (e.g., < 300). We present a method to detect inattention periods on a trial-by-trial basis to improve threshold estimates in psychophysical experiments using the adaptive sampling procedure Parameter Estimation by Sequential Testing (PEST). The performance of the algorithm was evaluated in computer simulations modeling inattention, and tested in a behavioral experiment on proprioceptive difference threshold assessment in 20 stroke patients, a population where attention deficits are likely to be present. Simulations showed that estimation errors could be reduced by up to 77% for inattentive subjects, even in sequences with less than 100 trials. In the behavioral data, inattention was detected in 14% of assessments, and applying the proposed algorithm resulted in reduced test-retest variability in 73% of these corrected assessments pairs. The algorithm complements existing approaches and can be adapted to sampling procedures other than PEST. Besides being applicable *post-hoc*, it could also be used online to prevent collection of biased data. This could have important implications in assessment practice by shortening experiments and improving estimates.

## 1. Introduction

Psychophysical procedures are used in many fields to assess sensory perception thresholds of stimuli in different modalities (e.g., vision, hearing, proprioception, or tactile sensation). During experiments, the assumption is generally made that the neurological and psychological mechanisms underlying perception are stationary. However, this may not always be the case, as fatigue, attention, learning, or a change in decision criteria or response bias occurring during the experiment can alter perception (Leek et al., 1991; Hall, 1983; Watson, 1980; Doll et al., 2015; Cameron et al., 2002; Carrasco et al., 2004; Carrasco, 2006; Cohen and Maunsell, 2011). Attentional deficits may be a common consequence of many types of brain damage, such as stroke, traumatic brain injuries, or brain tumors (Bruhn and Parsons, 1971; van Zomeren and van den Burg, 1985; Tuhrim, 1993; Rinne et al., 2013). Furthermore, according to pedagogical literature, the attention span is also limited in neurologically intact humans (7–20 minutes) (Bligh, 1998; Petty, 2004; David and Dukette, 2009; Dent and Harden, 2013). Thus, depending on the assessment paradigm, inattention can be a significant confound. Subjects may be distracted during the assessment and forget about the intensity of previously presented stimuli, or completely miss them, resulting in biased or random answers and thus incorrect outcome measures.

It has been shown that threshold estimates from adaptive stimulus placement procedures may be robust against large symmetric stimulus sequence excursions when sampling a psychometric function (Green et al., 1989). However, it is improbable that sustained periods of attention loss have a symmetric influence, but rather a detrimental effect on the mean and variance of the threshold estimates (Green, 1995). Furthermore, other parameters of the psychometric function such as the slope will also be affected (Leek et al., 1991). Finally, sustained inattention can lead to divergence of adaptive psychophysical procedures (refer to Treutwein (1995) and Leek (2001) for reviews), resulting in longer experiments, which in turn can be unfavorable for sustained attention. Thus, at the stimulus levels towards which the adaptive procedure may be converging and at which the most data is sampled, the psychometric function may be least stationary. In all cases, whether the estimate of the threshold is based on the final trials of an adaptive sequence (Taylor and Douglas Creelman, 1967), or on fitting a psychometric function to the proportions of correct responses for different stimulus levels (Treutwein and Strasburger, 1999), the resulting estimate may be unreliable and biased (Fründ et al., 2011), especially in 2-alternative forced-choice (2AFC) tasks (Green, 1995). Thus, a general challenge remains: How can biases resulting from inattention during psychophysical experiments be corrected for?

While inspection of recorded stimulus level sequences from adaptive procedures sometimes reveals a course of performance being visibly influenced by inattention, as the performance level suddenly decreases for a certain period, objective methods are required to detect such phases. Physiological signals such as electrodermal activity (EDA) could potentially be used to detect inattention intervals, as arousal has been found to be a strong predictor for attention (Prokasy and Raskin, 1973). However, depending on the situation causing inattention to the psychophysical task, it may not be obvious to interpret a change in the EDA signal, as the subject could be simultaneously inattentive to the task but aroused by a distractive situation, or as EDA could also respond to cognitive load. Furthermore, the measurement of EDA requires additional equipment and may not be applicable in some settings. Therefore, methods based solely on the stimulus sequence and responses recorded during the experiment would be of great advantage.

There have been only few attempts aiming at identifying sequences of questionable validity, as they may have been affected by inattention, and describing the overall stability of the threshold (Hall, 1983; Leek et al., 1991). However, these methods require computer simulations to compute normative tolerance limits and confidence intervals. As these statistical values are case specific (i.e., dependent on psychometric functions, sampling procedures, step sizes, etc.), new simulations would have to be performed for each specific experiment at hand, limiting the practicability of these approaches. This may motivate the use of empirically determined tolerance intervals (based on knowledge from prior experiments) (Hall, 1983). Together with the fact that they cannot identify specific trials affected by inattention within the sequences, nor correct for threshold estimate biases, these can only be used as preliminary screening tools. A different approach to improve parameter estimates of the psychometric function is to account for lapses (i.e., stimulus-independent errors, which can be caused by inattention) (Wichmann and Hill, 2001), or to improve sampling strategies of psychophysical procedures (Shen and Richards, 2012; Prins, 2013). While these approaches constitute viable solutions to identify questionable sequences as a whole, or to improve the goodness of fit and reduce variance of estimates in case of few isolated scattered lapses, these methods do not allow to detect local changes of attention on a trial-by-trial basis within a sequence.

A way to track threshold changes (i.e., varying threshold estimates throughout the experiment) consists in estimating the parameters of the psychometric function based on a subset of sequence data using a moving-window approach. This method has shown promising results in computer simulations using windows spanning 25 trials for modeled non-stationary thresholds following linear and exponential changes (Doll et al., 2015). Together with a classification or weighting algorithm, such a threshold tracking method could be used to identify and remove trials affected by attention loss. However, while this tracking method may be suitable for nociceptive detection thresholds (Doll et al., 2015), its applicability for assessments in other modalities may be limited for several reasons. First, the reliability of estimates may not be high enough even when using the complete sequence, as more than hundred trials may be required for acceptable confidence limits (McKee et al., 1985), which is especially the case for the 2AFC paradigm (King-Smith et al., 1994) and psychometric functions with shallow slopes (Green, 1995). Second, efficient adaptive procedures targeting different “sweet points” for the optimized estimation of parameters (e.g., Parameter Estimation by Sequential Testing (PEST) (Taylor and Douglas Creelman, 1967) or the maximum-likelihood procedure proposed by Shen and Richards (2012)) will not provide sufficient coverage of stimulus levels within each window given the limited window sizes. Fründ et al. (2011) proposed two approaches to adjust for the underestimation of parameter estimate confidence intervals (CIs) from non-stationary psychometric functions: correction of inference (by scaling the CIs) and correction of data (by estimating the number of independent trials and reducing each block of trials by the same number). While proving successful for a large number of trials, these approaches show limited or no improvement for relatively small numbers of trials (e.g., < 300). Moreover, this work focused on the CIs and not on the actual parameter estimates, and only symmetric non-stationarities were modeled. In summary, existing methods lack the ability to precisely identify periods of inattention within short sequences with a low number of trials to solve the underlying problem that a non-negligible amount of biased data from a non-stationary psychometric function may be included in the estimation.

We present a method to identify sustained inattention periods for improving threshold estimates by rejecting trials affected by non-stationarity. The proposed algorithm is based on data from the sequence of an adaptive sampling procedure and a stability criterion of the evolution of threshold estimates. Computer simulations were used to evaluate the correction method for models with different levels of simulated inattention. Additionally, behavioral data were used to illustrate and support the applicability of the proposed method as well as to validate the simulation and inattention model design. This work was framed around a scenario of a proprioceptive difference threshold assessment using PEST (Taylor and Douglas Creelman, 1967) in combination with a two-interval 2AFC assessment paradigm (Macmillan and Douglas Creelman, 2005).

## 2 Background

### 2.1 PEST: Parameter Estimation by Sequential Testing

The adaptive PEST algorithm (Taylor and Douglas Creelman, 1967) for assessing perception thresholds offers many advantages over other psychophysical procedures: its simple implementation, freely selectable convergence point (i.e., desired target of percentage of correct responses), and no required prior assumptions on the subject’s psychometric functions. PEST is based on a Wald sequential likelihood-ratio test (Wald, 1947), defining at each trial whether the stimulus level should remain at the same level or be decreased, respectively increased, by a specific step size, depending on the proportion of correct responses at the specific stimulus level. The stimulus level is maintained if *N*_*cor*_ ∈]*N*_*tot*_*×P*_*t*_ ± *W*[, where *N*_*cor*_ and *N*_*tot*_ correspond to the correct and total number of trials at the current stimulus level, *P*_*t*_ the desired target performance, and *W* the Wald sequential likelihood-ratio test parameter. Direction reversals (switching from decreasing to increasing steps, or vice versa) allow PEST to converge towards the threshold in an oscillatory manner. The next stimulus level to be presented is defined by a set of heuristic rules taking into account responses of past trials:

- The step size is halved at every direction reversal.
- The first and second step in the same direction are of same size.
- The third step is double the second if the size of the step immediately preceding the last reversal resulted from a doubling, or same otherwise.
- The fourth and additional steps in the same direction are the double of the preceding step size.

In a 2AFC paradigm, the desired target of percentage of correct responses *P*_*t*_ is 75%, and a Wald sequential likelihoodratio test parameter *W* of 1 is recommended by Taylor and Douglas Creelman (1967) for a reasonable trade-off between more precise estimates and longer assessments. PEST requires two starting parameters: the start level and the start step, which should be selected if possible using some prior knowledge about the distribution on thresholds to be assessed in an experiment. The PEST algorithm features different termination criteria: maximum total number of trials, maximum number of trials at the same stimulus level, and minimum step size. As soon as one of the termination criteria is fulfilled, the outcome measure (i.e., threshold) of the assessment is defined by the level of the next stimulus which would be presented.

PEST has often been used in psychoacoustic experiments using auditory stimulus levels in dB (Taylor and Douglas Creelman, 1967; Hall, 1981; Leek, 2001). However, it has been suggested to implement the PEST algorithm in a logarithmic domain for “non-logarithmic stimuli”, such as absolute angles in a proprioceptive assessment, to avoid inefficient behavior (e.g., due to zero crossings) (Rinderknecht et al., 2014). This adapted version making use of the logarithmic domain was used in the following computer simulations and behavioral experiment in stroke survivors on finger proprioception. As a two-interval 2AFC paradigm was used, the absolute angular difference between the two consecutive stimuli of one trial are referred to as stimulus level *x*, or in short “level”.

### 2.2 Fitting the psychometric function

Despite PEST being an adaptive algorithm able to directly provide an estimate of the threshold at convergence, some scenarios may not allow convergence of the algorithm, for example due to a premature termination of the experiment or too severe termination criteria. One robust solution to address this issue, consists in using all available data of the PEST sequence (i.e., pairs of stimulus level *x* and response) and estimating the threshold by fitting a psychometric function *ψ*(*x*) (Hall, 1981) using a Maximum Likelihood criterion (Prins and Kingdom, 2009). This approach has been validated in a previous experimental study (Rinderknecht et al., 2014). Therefore, in both simulations and behavioral experiments of the present work, the thresholds were estimated based on the fitting procedure.

The psychometric functions *ψ*(*x*) are defined by the proportion of correct responses at different stimulus levels *x*:

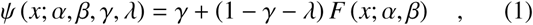

where the generic sigmoid function *F*(*x*; *α*, *β*) was chosen to be a cumulative normal function *F*_*Gauss*_(*x*; *μ*, *σ*) with a mean *μ*, corresponding to the inflection point of *Ψ*(*x*), not necessarily being the same as the generic threshold parameter *α*, and standard deviation *σ*, inversely proportional to the slope parameter *β*:

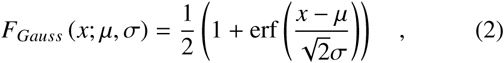

erf(*x*) being the standard definition of the error function:

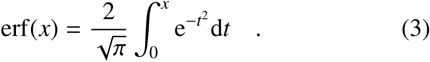

Given the 2AFC paradigm used in this proprioceptive assessment framework, the guessing rate *γ* was set to 0.5. When fitting simulated and behavioral data in the present work, the lapse rate *λ* was allowed to vary ∈ [0, 0.1] to reduce estimation bias and account for isolated scattered lapses (Wichmann and Hill, 2001). Note that the interval was selected to be slightly larger than [0, 0.06], as used in the work by Wichmann and Hill (2001), in order to account for a potentially higher probability of inattention in stroke patients (Tuhrim, 1993; Rinne et al., 2013). If *λ ≠* 0, the threshold does not correspond to the inflection point *μ* anymore. Consequently, the threshold *x*_*T*_ is always defined at a 75% performance level: *x*_*T*_ = *Ψ*^−1^(0.75). Furthermore, following the suggestions from Strasburger (2001) to obtain comparable values across different studies, the actual slope is defined as:

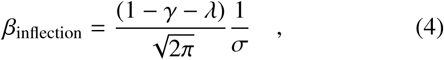

being the first order derivative of *Ψ*(*x*) at the inflexion point. At the same time, this slope corresponds to the maximal slope of the sigmoid function. **Figure 1** illustrates the different functions and definitions of the various parameters.

**Figure 1.**
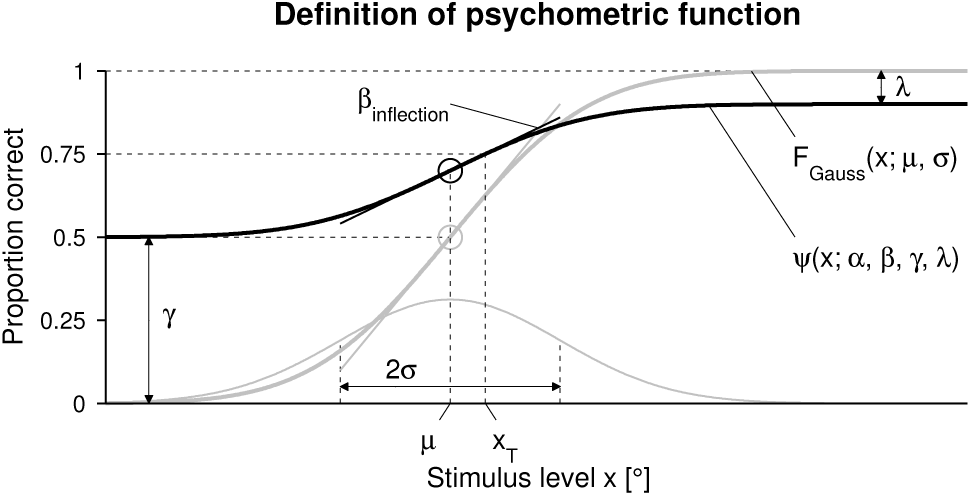
Definition and parameters. Definition and parameters of the psychometric function *Ψ*(*x*; *α*, *β*, *γ*, *λ*) (bold black sigmoid), the cumulative normal function *F*_*Gauss*_(*x*; *μ*, *σ*) (bold gray sigmoid), and its underlying normal probability density function (gray). The inflection points and slopes at these points are indicated by circles and lines, respectively, in the colors of the sigmoids.

## 3 Method to correct for sustained inattention

The aim of the proposed method to correct for sustained inattention consists in identifying the period in which the subject is inattentive, and excluding the biased data collected during this period when using the fitting process for threshold estimation. Thus, the time point at which the subject’s performance starts to decrease due to inattentiveness and the potential time point at which the performance starts improving again (i.e., subject being attentive again) have to be estimated. We propose to analyze both the PEST sequence and the evolution of threshold estimates along the psychophysical assessment.

The inattention detection method is based on heuristic rules. Using visual inspection of samples from previously recorded behavioral data (different proprioceptive assessments in healthy subjects) as well as preliminary simulations, the method’s face validity was tested and its parameters were optimized.

Initially, the first trial which is followed by four consecutive PEST level increases without reversal (denoted as *t*_PEST,4up_, *t* standing for trial) is identified. If this trial is within the first 8 trials, it is disregarded and replaced by the next occurrence. This allows the PEST algorithm to adapt the stimulus level in initial trials if the start level was chosen too low. Note that this parameter strongly depends on the choice of the start level and start step, and tuning would require detailed knowledge of the threshold distribution of the subject population in question. However, in general, the start level should be chosen high enough to make the task easily understandable for the subject and thus render this additional rule superfluous.

Subsequently, similar to the sequence stability idea of Leek et al. (1991), the stability of the evolution of the threshold estimate is calculated by defining adaptive boundaries. By fitting the psychometric function to the data (i.e., proportion of correct responses at presented stimulus levels) available until trial *t*, the evolution of the threshold estimate 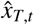 is calculated (**Figure 2 (Top)**). This threshold estimate is smoothed 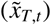 with a moving average filter (span of 7 trials, based on optimization) to remove spiky estimation errors, which can occur when only a small number of data points are available during the fitting process. To deal with the endpoint, the array of threshold estimates is padded (by replicating the last available value 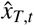 for 3 trials) prior to the smoothing process and truncated to the original length after smoothing. When using the method offline, the last available value 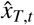 corresponds to the last trial of the experiment 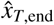, whereas when using the method online, this smoothing procedure is repeated on the last few trials, as the four previous trials to the current trial 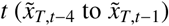 will change due to the moving padding because new data is available. The mean 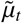 and standard deviation 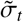 of the smoothed threshold estimate evolution 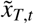 are calculated The mean for each trial *t* according to the following equations:

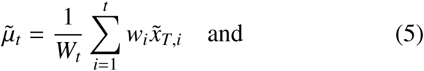

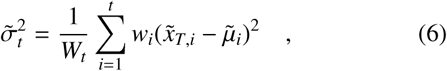

using exponential weights *w*_*i*_ for the trials 1 to *t* with a forgetting rate *ε* = 0.5:

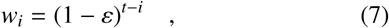

normalized with *W*_*t*_ being the sum of all weights *w*_*i*_ up to trial *t*:

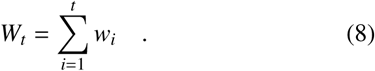

**Figure 2.**
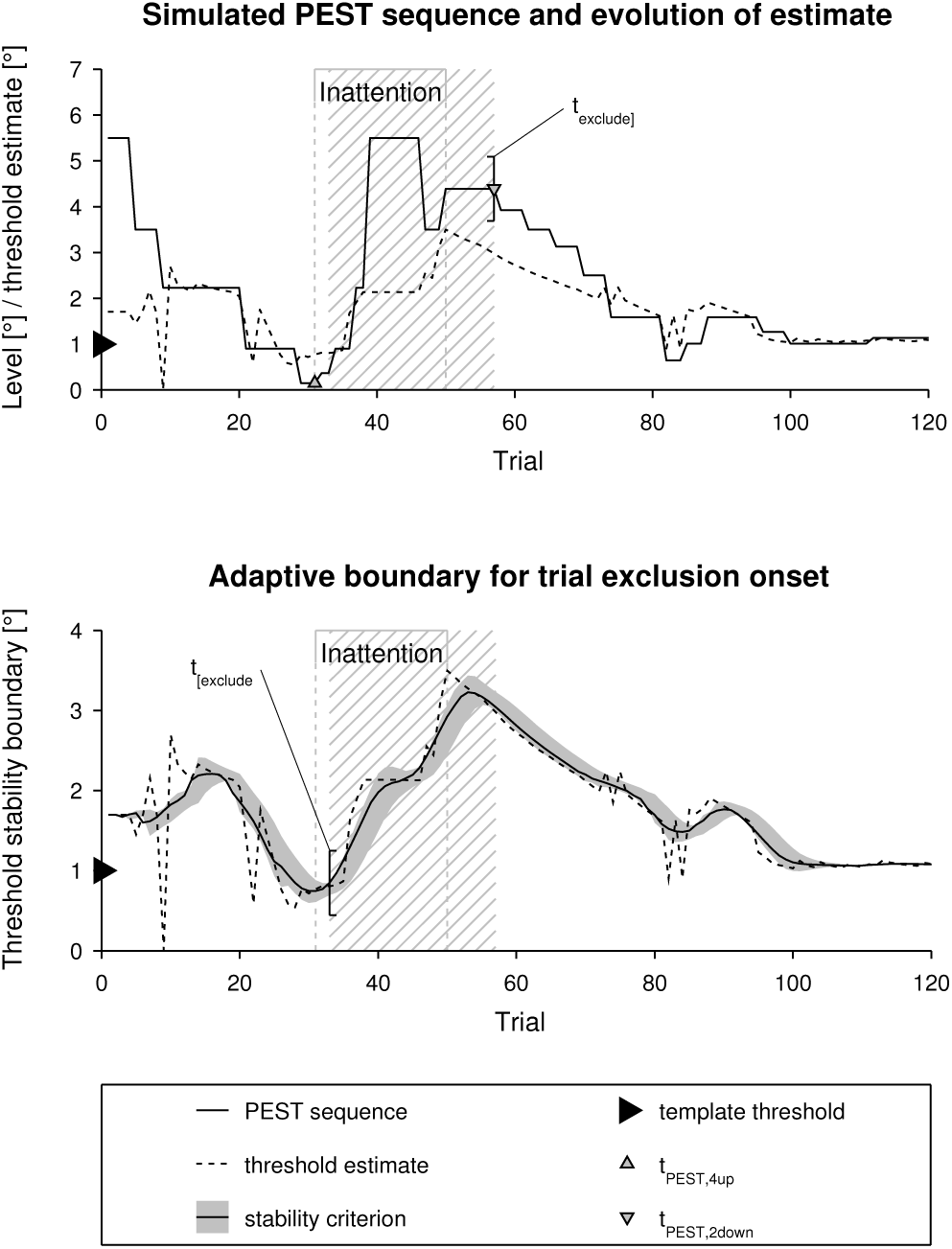
Example of a PEST sequence and corresponding adaptive boundary for inattention correction in a simulated run for template 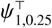 and *λ*_inattention_ = 0.5. (Top) Level of the PEST sequence (solid black line) and evolution of the threshold estimate 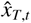 (dashed black line). The upward-pointing gray triangle designates *t*_PEST,4up_, the first trial being followed by four consecutive level increases without reversal, and the downward-pointing gray triangle designates *t*_PEST,2down_, the first (after detected inattention onset) trial followed by two consecutive level decreases without reversal. The period of trials which are excluded by the inattention correction method are indicated by the hatched region spanning from *t*_[exclude_ to *t*_exclude]_. The modeled inattention period is labeled and indicated by the gray bracket and gray dashed lines, and the threshold of the template by the black triangle. **(Bottom)** Raw threshold estimate 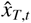 (dashed line), exponentially weighted, smoothed threshold estimate average 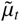 (solid line), and threshold stability boundary based on 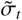 (gray shaded band).

By using this exponential weighting, more recent values are taken into account more strongly and 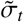 adapts its size depending on how stable the estimate 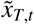 **Figure 2 (Bottom)** This can be used to define a boundary to detect a significant change in the trend of the estimate: Starting with the first occurrence after trial *t*_PEST,4up_ where the following condition 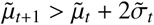 is met (defined as trial *t*_[exclude_, with symbol [ denoting the start of exclusion), the trials are excluded for the calculation of the final threshold estimate. The range 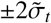 corresponds to about 95% of the data and has shown to be a reasonable choice in preliminary simulations.

Analogous to *t*_PEST,4up_, the first trial followed by two consecutive PEST level decreases (8 correct responses would lead to 2 level decreases, considering *P*_*t*_ = 75% and *W* = 1) after *t*_[exclude_ is identified and denoted as *t*_PEST,2down_. This trial *t*_PEST,2down_ simultaneously corresponds to the last trial to be excluded (*t*_exclude]_, where symbol] denotes the end of exclusion). All following trials (including the 2 level decreases) are reconsidered for the calculation of the final threshold estimate and all trials from *t*_[exclude_ to *t*_exclude]_ are excluded.

## 4 Computer simulations

Computer simulations are a valuable tool to evaluate the ability of the proposed method to correct for sustained inattention, as when modeling a subject, its psychometric function (also referred to as template) and modeled inattention are completely defined. Furthermore, today’s computational power offers the possibility to simulate an extensive number of experimental runs to calculate a precise distribution of estimation errors.

Computer simulations were run with the following aims: (i)evaluate the performance of inattention correction in templates featuring different levels of modeled inattention, and (ii)validate that the proposed method does not significantly affect threshold estimates in absence of inattention. Thus, in the latter, the same templates were used, with inattention correction but without modeled inattention. The same simulation without inattention model but without applying the inattention correction method served as a baseline for the validation. To obtain representative results from the computer simulations (see, e.g., Kollmeier et al. (1988)), simulated templates and parameters of the sampling procedure were selected based on our prior experimental knowledge and experience.

### 4.1 Methods

#### 4.1.1 Templates

Twelve different subject templates covering a difference threshold space (*x*_*T*_: ∈ {1, 2, 4 }°) and a slope space (*β*_inflection_: ∈{0.0625, 0.25, 0.5, 1 }/°) were simulated. These values are representative for proprioceptive perception in healthy adults (Choi et al., 1995; Fry-Welch et al., 2003; Brewer et al., 2005; Tan et al., 2007; Lambercy et al., 2011; Rinderknecht et al., 2014; Elangovan et al., 2014; Cappello et al., 2015). The guess rate *γ* was 0.5 and the lapse rate *λ* was 0 for all templates. In the following text, the short notation 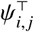 (symbol 𝒯 denotes a template) is used for templates, with index *i* denoting the threshold space parameter value, and index *j* the slope space parameter value (*e.g.*, 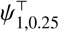 for *x*_*T*_ = 1^°^ and *β*_inflection_ = 0.25/°).

#### 4.1.2 Inattention model

Inattention can be modeled in different ways: by changing the threshold (Hall, 1983; Leek et al., 1991; Cameron et al., 2002; Doll et al., 2015), the slope (Cameron et al., 2002), the lapse rate (Green, 1995; Wichmann and Hill, 2001), or a combination thereof (Cameron et al., 2002; Fründ et al., 2011). Depending on the concrete implementation of the psychometric function, modifying function-specific parameters may have different effects on the psychometric function (e.g., in the implementation by Green (1995) adding a lapse rate > 0 decreases the actual slope of the psychometric function without changing the slope parameter).

As the slope parameter of the psychometric function describes the reliability of stimulus detection or discrimination (Strasburger, 2001), it seems appropriate to decrease the slope *β* of the psychometric function to model inattention. However, this would lead to higher performance for stimulus levels below the inflection point, which is contradictory to the fact that performance is assumed to decrease if the subject is inattentive. Consequently, the threshold parameter *α* would need to be increased to resolve this issue. While it also seems plausible that loss of attention would result in a higher number of lapses, the lapse rate *λ* could also be increased. As a matter of fact, increasing *λ* in our implementation decreases the actual slope *β*_inflection_ (see Equation 4) and increases the actual threshold *x*_*T*_ = *Ψ*^−1^(0.75). Thus, the performance of the resulting proprioceptive function is lower compared to the performance of the original psychometric function, for all stimuli *x* > 0. Hence, we hypothesize that changing the lapse rate is the most suitable approach, which is in addition simpler than changing multiple parameters simultaneously.

A set of different lapse rates were simulated (*λ*_inattention_ ∈{0, 0.25, 0.5 }/°). While *λ*_inattention_ = 0.5 corresponds to complete inattention (i.e., guessing) and *λ*_inattention_ = 0.25 leads to a psychometric function with asymptotic value of 0.75, *λ*_inattention_ = 0 models a perfectly attentive subject.

#### 4.1.3 Procedure

The following parameters were used for the PEST procedure: start level of 5.5°, start step of 2°, desired performance level *P*_*t*_ of 75%, and Wald sequential likelihood-ratio test parameter *W* of 1. These parameters proved successful in prior experiments assessing proprioceptive difference thresholds of finger joints (Rinderknecht et al., 2014). Each simulated experiment lasted for 120 trials and no other PEST termination rule was used. Observations from previous experiments (e.g., (Rinderknecht et al., 2014; Metzger et al., 2014) with healthy subjects and stroke patients) and ongoing proprioceptive assessments using PEST conducted by our group described subjects losing attention after 5–10 minutes of performing the task (corresponding to about 20–40 trials). This time interval is similar to the widely used claim in teaching guides that student’s attention span ranges from 7–20 minutes (Bligh, 1998; Petty, 2004; David and Dukette, 2009; Dent and Harden, 2013). However, this claim is mostly based on subjective observations and is supported by little well-controlled studies (Wilson and Korn, 2007). Nevertheless, it would be reasonable that the attention span in our 2AFC task is reduced, as the task is very repetitive and active participation is limited because the subjects receive only passive stimulation. Therefore, the onset of the modeled inattention was empirically chosen to follow a normal probability distribution centered around trial *t* = 30 with a standard deviation of 10 trials: **𝒩** (30, 10). The sustained inattention period was modeled for 20 consecutive trials. Since the performance of the inattention correction method will be evaluated as a function of trials ∈ [1, 120], the upper bound of 120 trials only presents a limit for analyzing the evolution of the performance and should be sufficiently high to show its asymptotic behavior. Furthermore, based on practical experience in our group, it can be noted that 120 trials are towards the limit of practical feasibility in proprioceptive assessments using the two-interval 2AFC paradigm, as (i) the presentation of passive movements is time-consuming, and (ii) the task becomes cognitively very demanding and tiring when presented stimuli are around the perception threshold. In a clinical setting, a maximum of 60 trials corresponding to 15 minutes might be more realistic, as assessments are required to be quick to administer (Gresham et al., 1996). For each inattention model, the experiment was simulated 1000 times per template 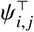, leading to a total of 36000 simulated experimental runs 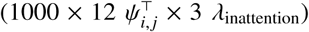 All simulations were performed in MATLAB R2014a (MathWorks, Natick, MA, USA).

#### 4.1.4 Data analysis

For all simulated templates 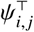 and modeled inattention levels, the absolute error (AE), constant error (CE), and variable error (VE) were calculated according to the standard implementations (Schmidt and Lee, 2011) to evaluate the estimation performance in both cases, without and with inattention correction. All threshold estimates larger than 20^°^ were considered as outliers, as such large values are unreasonable in this specific scenario of finger proprioception. However, it is important to not disregard such poor estimates, as they affect the general quality of a psychophysical assessment. Thus, they were included when evaluating the inattention correction method. To reduce their severe effect on error metric statistics, threshold estimates were saturated to a ceiling value of 20°. The average mean ± SD in the individual evaluation metrics and the absolute and percentage improvements were calculated across all templates 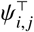 for each inattention model without and with inattention correction. In order to calculate the average bias for the performance evaluation across the different templates 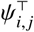, the sign was removed from each CE of each template 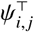 by taking the absolute value (|CE|). Positive improvement represents either reduction of variance or bias. Furthermore, the exclusion frequency for each trial and the distributions of trials at which the correction method starts excluding the data (*t*_[exclude_) and trials at which data is reconsidered after the detected inattention period (*t*_exclude]_) were calculated. Before computing mean ± SD, PEST sequences were synchronized *post-hoc* to the same modeled inattention onset, as the inattention onset was randomly distributed across the simulated runs. Additionally, percentages of sequences being truncated before, during or after modeled inattention, or not at all, were computed. All analyses were conducted in MATLAB R2014a.

### 4.2 Results

An example of inattention detection and trial exclusion in a simulated run for template 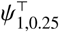 and *λ*_inattention_ = 0.5 is presented in **Figure 2**. This example clearly illustrates how the PEST sequence diverges rapidly after modeled inattention onset (**Figure 2 (Top)**). After the simulated inattention onset at trial 31, the performance dictated by the altered psychometric function changes for the presented level, and the following ratio of correct versus incorrect responses drops. This leads to the subsequent increases of the level and influences the threshold estimate 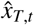 in a similar way (**Figure 2 (Top)**). Despite the last trial of the modeled inattention being at 50, the last trial of exclusion *t*_exclude]_ is at 57, before the PEST sequence starts decreasing and converges towards the true threshold. Data from the plateau before the level decrease is always excluded for the threshold estimate, as the first few trials of the plateau may be still partly influenced by inattention. The onset of data exclusion *t*_[exclude_ is defined based on the stability of the threshold estimate. The evolution of the threshold estimate 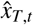, the exponentially weighted, smoothed threshold estimate average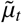, and the threshold stability boundary based on 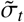 are illustrated in **Figure 2 (Bottom)**. This plot visualizes how the width of the stability boundary becomes wider the more the threshold estimate varies (e.g., shortly after inattention onset) and the narrower the more stable it is (e.g., towards the end of the sequence). Thus, the rapid change in 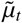 originating from inattention can be detected shortly after inattention onset (*t*_[exclude_ = 33). Furthermore, this example shows spiky estimation errors in 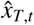, predominantly towards the beginning of the sequence, because of which it is necessary to use the smoothed 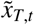 for the inattention detection. The resulting psychometric functions (without and with inattention correction) after 60 and 120 trials are shown in **Figure 3** together with the template 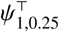. In both plots it can be seen that, in the condition where no inattention correction is applied, the proportions of correct responses for the same levels is lower (e.g., at 2.2°), and that there is additional data with low performance at above-threshold levels (e.g., at 4.4°). This results in a downwards shift of the psychometric function creating a rightwards shift (i.e., increase) of the threshold estimate. Furthermore, it should be noted that in this specific example the estimated psychometric function without inattention correction matches the general shape (among others the actual slope and the lapse rate) of the template 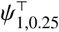 better than with inattention correction. The large lapse rate of the fit with inattention correction may seem counterintuitive. It may be caused mainly because of the interaction between the high performance at low level 0.90^°^ and the proportion of correct responses at 2.22^°^ which is lower than the psychometric function of the template. Nevertheless, correction for inattention improves the threshold estimate, especially if the maximum number of trials of an assessment is short (e.g., 60 trials, **Figure 3 (Top)**). The longer the assessment, or the more trials follow after the inattention period, the more the two estimates approach each other (e.g., 120 trials, **Figure 3 (Bottom)**).

**Figure 3.**
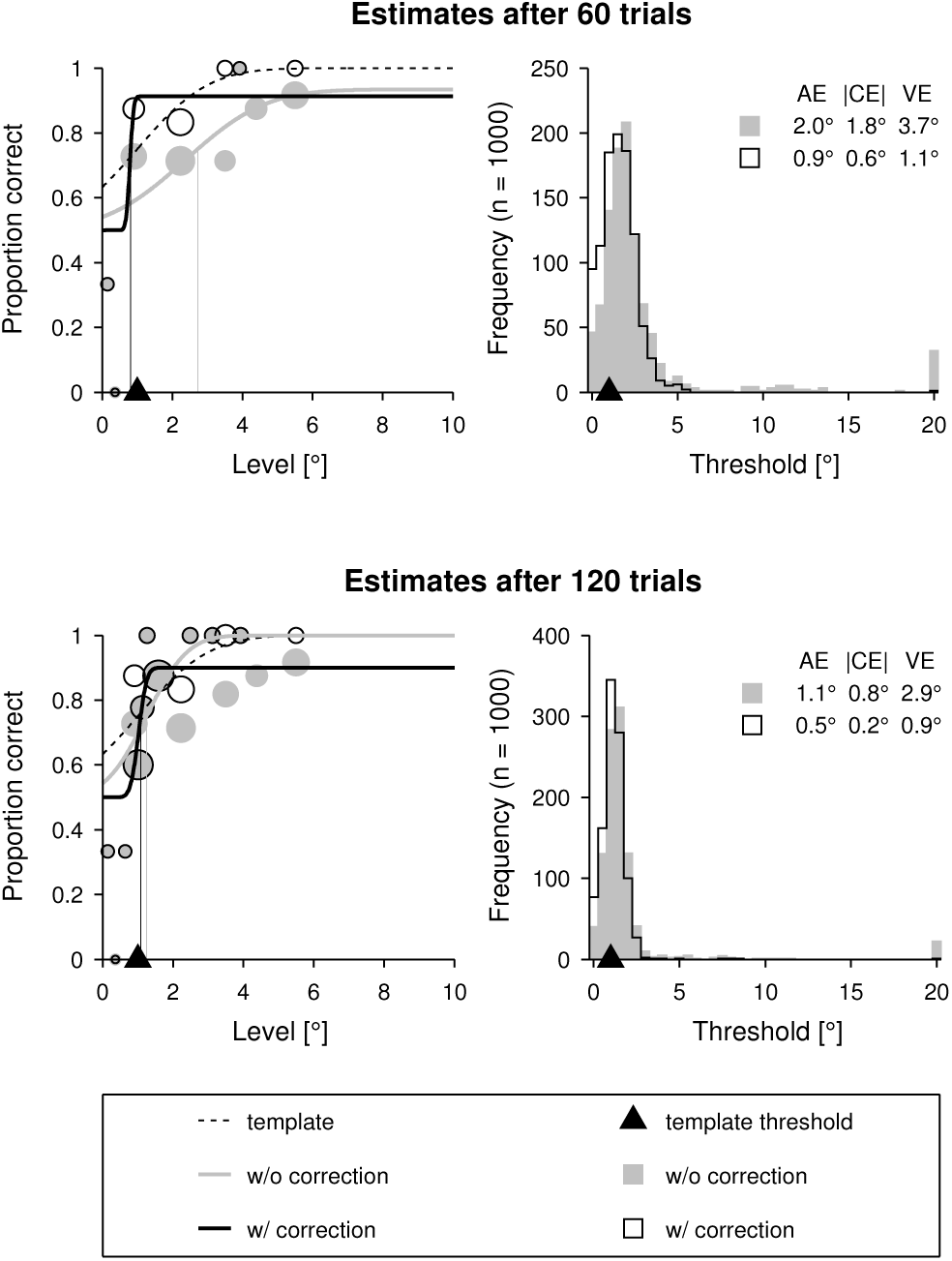
Example of fitted psychometric functions and threshold estimate distributions without and with inattention correction. Example of fitted psychometric functions without and with inattention correction from a single simulated run for template 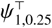 (for the same run as in **Figure 2**) **(Left)** and threshold estimate distributions (bin resolution of 0.5°) from 1000 simulated runs of the same template **(Right)**, after 60 **(Top)** and 120 trials **(Bottom)**, with *λ*_inattention_ = 0.5. **(Left)** Psychometric function of the template 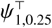 (dashed black sigmoid), estimated psychometric function without inattention correction (solid gray sigmoid), and estimated psychometric function with inattention correction (solid black sigmoid) after 60 trials. The thresholds of the two latter psychometric functions are indicated by vertical dashed lines in the respective colors. The proportions of correct responses are indicated as circles (gray: without inattention correction, black outlines: with inattention correction) for the different presented levels. The diameter is proportional to the number of presentations. The threshold of the template is indicated by the black triangle. **(Right)** Comparison of threshold estimate distributions (bin resolution of 0.5°) without (gray fill) and with (black outline) inattention correction. The threshold of the template 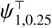 is indicated by the black triangle. Reported are the absolute error (AE), absolute value of the constant error (CE), and variable error (VE) of the estimate distributions.

The estimation errors for the different inattention models (i.e., different *λ*_inattention_) for the conditions with and without inattention correction are shown in **Figure 4**. The plots show that the estimation performance was severely affected by large *λ*_inattention_, reaching errors of the same order of magnitude as the thresholds to be estimated, when inattention was not corrected for. By applying the method to correct for sustained inattention, these estimation errors could be reduced. However, the they were not eliminated completely, and remained larger than in the optimal case, where perfect attention (*λ*_inattention_ = 0) was modeled and no correction was performed. The largest estimation errors were found around trial 50, which corresponds to the average of the distribution of the end of the inattention period (center of the normal random distribution 𝒩(30, 10) of the modeled inattention onset plus20 trials of sustained inattention). After this peak, the estimate errors decreased exponentially with the amount of additional data collected after the inattention period, independently of whether inattention was corrected for or not. Large oscillations in estimation performance during the first 20 trials result from the low number of data points (mostly at high levels with high performance rates) available for fitting the psychometric function.

**Figure 4.**
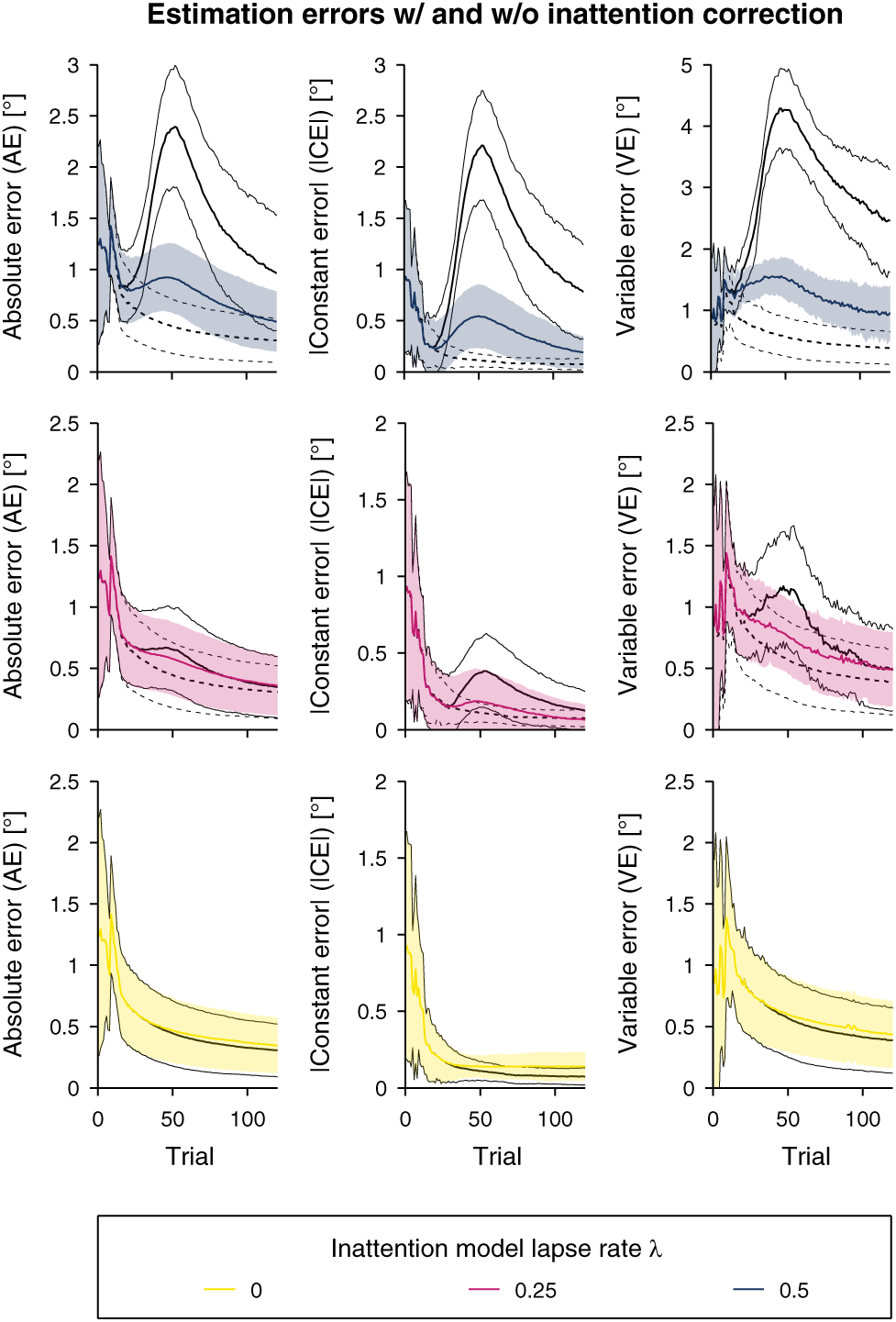
Estimation errors for the different inattention lapse rates. Estimation errors (mean ± SD of the absolute error (AE), absolute value of the constant error (|CE|), and variable error (VE) across the different templates 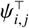) for the different inattention lapse rates *λ*_inattention_. The colored lines and shaded bands indicate the estimation errors after inattention correction for different *λ*_inattention_ (**(Top)** *λ*_inattention_ = 0.5, **(Middle)** *λ*_inattention_ = 0.25, **(Bottom)** *λ*_inattention_ = 0). For the same inattention models, the estimation errors without correction are represented by solid black lines. As a reference, the estimation errors are visualized for no modeled inattention (i.e., perfectly attentive subject) with dashed black lines. Note that for *λ*_inattention_ = 0, the two latter are identical.

The frequency of threshold estimates ceiling at 20^°^ could be reduced by applying the inattention correction, as presented in **Figure 5**. For *λ*_inattention_ = 0 and *λ*_inattention_ = 0.25, it was below 0.1% independent of the number of trials, and below 0.4% on average across different templates 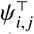 for *λ*_inattention_= 0.5, when inattention was corrected for. In the condition where the inattention correction method was not applied, the frequency of ceiling effects was consistently higher, especially for higher inattention lapse rates (e.g., reaching up to 5% on average for *λ*_inattention_ = 0.5 for a sequence length of 50 trials).

**Figure 5.**
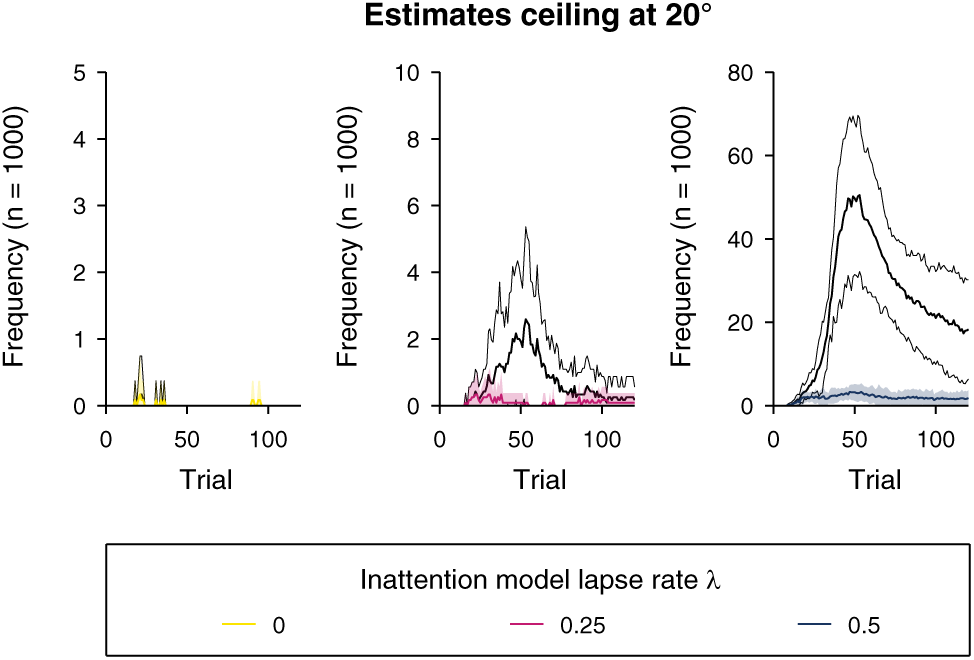
Frequency of ceiling threshold estimates for the different inattention lapse rates. Frequency of ceiling threshold estimates (mean ± SD across the different templates 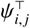) for the different inattention lapse rates *λ*_inattention_ (**(Left)** *λ*_inattention_ = 0, **(Middle)** *λ*_inattention_ = 0.25, **(Right)** *λ*_inattention_ = 0.5). The colored lines and shaded bands indicate the frequency of ceiling effects with inattention correction for different *λ*_inattention_, and the solid black lines without inattention correction.

The relative and absolute improvements with inattention correction in AE, |CE|, and VE are visualized in **Figure 6** for different *λ*_inattention_. For complete inattention (*λ*_inattention_ = 0.5), there was an estimation performance improvement of up to 62%, 77%, and 66% in the error metrics when correcting for inattention (maximum absolute improvement: 1.5^°^ for AE, 1.7^°^ for |CE|, and 2.8^°^ for VE). Estimation performance decreased for all error metrics in the case of absence of inattention (*λ*_inattention_ = 0). However, when inspecting the absolute improvements (**Figure 6 (Bottom)**), this performance decrease of up to 0.06^°^ for AE, |CE|, and VE was marginal. The large oscillations and negative peaks in the relative improvement (mainly for |CE|) originate from the division by constant errors near zero and were not representative for all templates 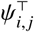

**Figure 6.**
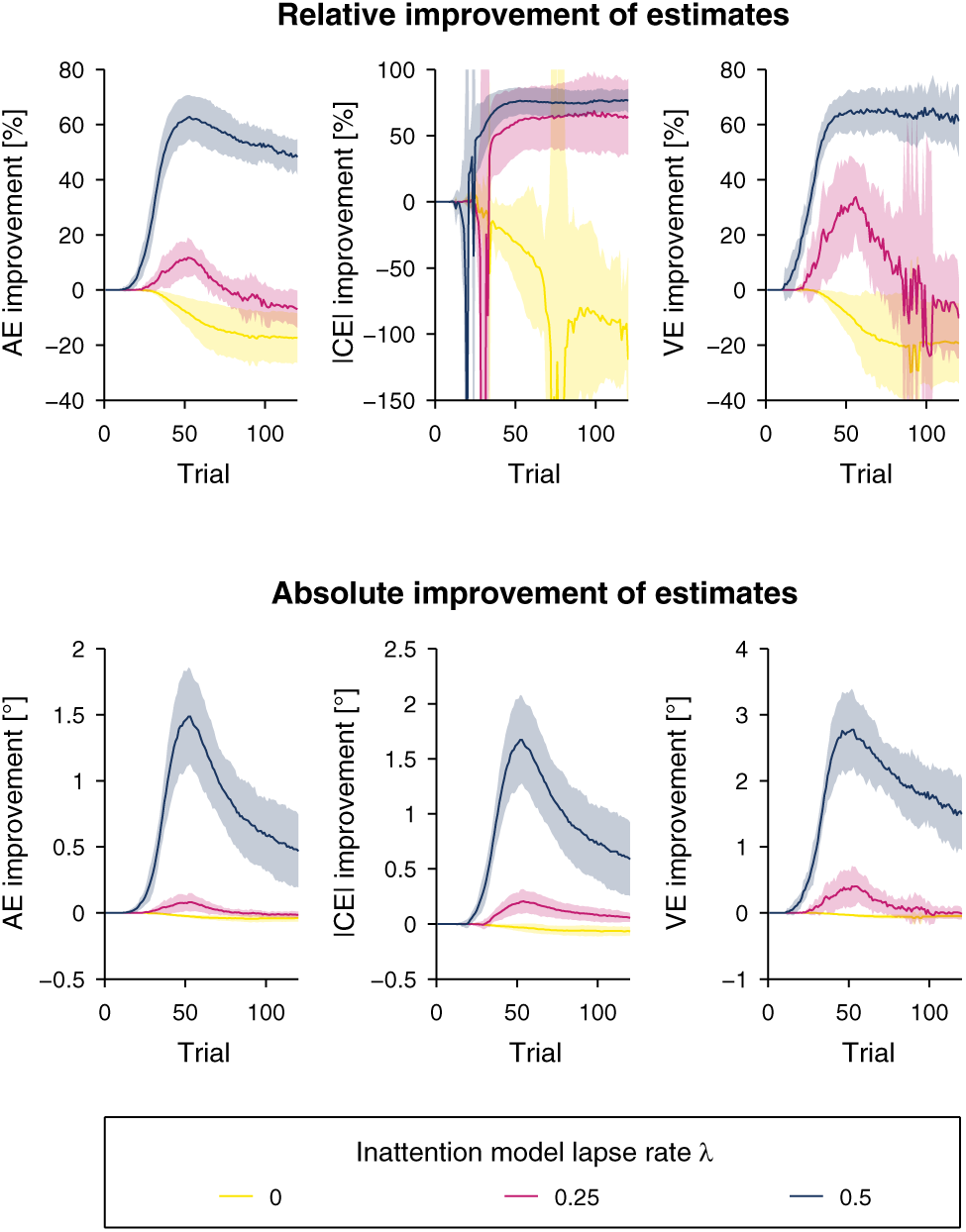
Improvement of the threshold estimation quality when using the inattention correction method vs. when no correction for inattention is performed. (Top) Relative improvement (mean ± SD across the different templates 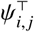) of the absolute error (AE), absolute value of the constant error (|CE|), and variable error (VE) for the different inattention lapse rates *λ*_inattention_. **(Bottom)** Absolute improvement of AE, |CE|, and VE for the different *λ*_inattention_.

A summary of the distribution of excluded trials (with *post-hoc* synchronized inattention onsets), as well as exclusion onset (*t*_[exclude_) and exclusion end (*t*_exclude]_) is presented in **Figure 7**. It can be seen that for *λ*_inattention_ = 0.5 and *λ*_inattention_ = 0.25, trials around the end of the inattention period were excluded based on the correction method in about 51% and 17% of the simulated runs, respectively. It is also noteworthy that the frequency at which trials were excluded increased linearly for trials during the inattention period and decreased even more rapidly around 2 trials after the end of the modeled inattention period. In the condition without inattention (*λ*_inattention_ = 0) the frequency of exclusion increases steadily with increasing trial number until around trial 70 after “theoretical inattention onset”, but does not reach more than 13%. The distribution of detected inattention onset and end (**Figure 7 (Middle)**) corresponds to the derivative of the distribution of excluded trials (**Figure 7 (Top)**) and highlights the aforementioned properties. This plot also shows that there was a small number of simulated runs (around 0.3% for each trial) where inattention is erroneously detected outside (i.e., after) the modeled inattention period.

**Figure 7.**
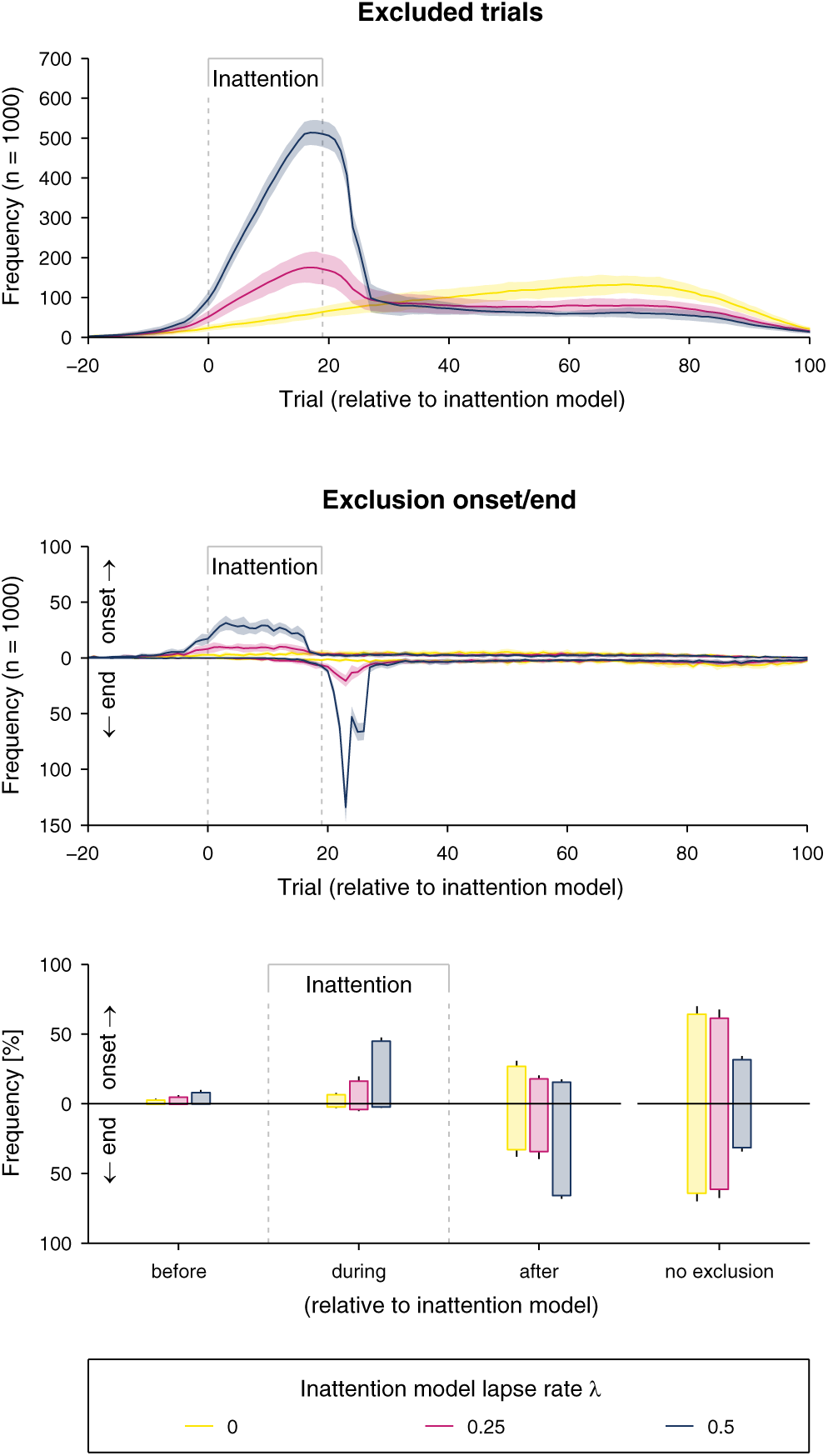
Distributions of data exclusion based on the inattention correction method with *post-hoc* synchronized inattention period. (Top) Frequency with which each trial is excluded. Reported are mean ± SD across the different templates 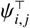 for the different inattention lapse rates *λ*_inattention_. The modeled inattention period is labeled and indicated by the gray bracket and gray dashed lines. **(Middle)** Distributions of exclusion onset (*t*_[exclude_) and exclusion end (*t*_exclude]_). Note that the distribution of the exclusion end is visualized on the inverted axis to make start and end distributions better distinguishable on the same plot. **(Bottom)** Grouped frequency (in per cent) of the exclusion onset and end occurrence (*before*, *during*, or *after* inattention period, or *no exclusion* if the correction method did not exclude any trials). For consistency, the exclusion end is again visualized on the inverted axis. Reported are mean ± SD.

The number of onsets (*t*_[exclude_) and exclusion ends (*t*_exclude]_) *before*, *during*, or *after* the modeled inattention period, or where *no exclusion* occurred, are summarized in **Figure 7**. The inattention onset during the inattention period increased with higher inattention, and was detected in 45% of the simulated runs for *λ*_inattention_ = 0.5 and 16% for *λ*_inattention_ = 0.25 during the inattention period. For all inattention models, the inattention onset was falsely detected before the actual onset of inattention in less than 8% of all simulation runs. The frequency of cases where no exclusion occurred decreased with higher inattention lapse rates, and was 64% for *λ*_inattention_ = 0, 61% for *λ*_inattention_ = 0.25, and 32% for *λ*_inattention_ = 0.5.

### 4.3 Discussion

Through the computer simulations, we aimed to (i) investigate whether and by how much the proposed correction method for sustained inattention improves the threshold estimates in the presence of sustained inattention, and (ii) to validate that the inattention correction does not significantly bias the threshold estimates in absence of sustained inattention. The main results demonstrated that estimation errors could be reduced by up to 77% in case of complete inattention, and that in case of absence of inattention the errors were only marginally different from the condition where no inattention correction was applied. Furthermore, when correcting for inattention, there were very few poor estimates > 20°, compared to when inattention was not corrected for, in which case it occurred in up to 9% of the simulated runs for some templates.

The excluded trials were predominantly within the modeled period of inattention, and the end of the simulated inattention phase (*t*_exclude]_) could be well identified. The short lag of *t*_exclude]_ (resulting in overestimation of the inattention period) can be explained by the Wald sequential likelihood-ratio test, as a minimum ratio of correct versus incorrect responses are required for the PEST level to be decreased. Since part of the trials at the stimulus level towards the end of the modeled inattention are still influenced by inattention (i.e., lower performance), a certain number of additional correct trials may be required after the end of the modeled inattention period to reach the condition for the PEST level to change. Since psychophysical assessments are based on probabilities and are not a fully deterministic process, it is possible that there is a small number of PEST sequences showing temporary divergences similar to sustained inattention, but which do not originate from the inattention model. However, as demonstrated by the results of the simulations, erroneous detection of inattention (in up to 8% of the runs) is not detrimental to the estimation quality. This can be supported by the fact that, in the case of absence of inattention, the estimation performance is only marginally changed, considering all erroneous detections before, during, and after the modeled inattention period (estimation quality decreased by less than 0.1°, which is at least one order of magnitude smaller than the thresholds, see **Figure 7 (Bottom)** for *λ*_inattention_ = 0). It should be noted that the number of inattention onset detections after the end of the modeled inattention period depends on the number of simulated trials (in this case 120 trials), as there remains a small probability of false detection which is summed up over the trials. Therefore, the reported frequencies in **Figure 7** should be interpreted with care. It may seem counterintuitive that almost 32% of the simulated runs with *λ*_inattention_ = 0.5 and 61% with *λ*_inattention_ = 0.25 were not truncated despite the presence of modeled inattention. Yet, it should be considered that if the subject is distracted (i.e., forced to guess), this corresponds to a performance rate of 50%. As a consequence, the levels of a PEST sequence do not increase as rapidly as when all responses were incorrect, or as it would be the case in a yes–no detection or same–different discrimination paradigm. Therefore, the level increase may very well take more than 8 trials (and even more for *λ*_inattention_ = 0.25), and thus may not reach four level increases within the 20 trials with modeled inattention. Furthermore, since the simulation of responses with and without modeled inattention is based on a stochastic process, there can be cases where the PEST level is decreased within the increasing step series, due to “lucky” guesses. The proposed method may not identify such events as inattention but consider them as normal oscillatory fluctuation towards the threshold. Another reason could be that PEST did not converge sufficiently in the preceding trials and therefore the width of the threshold estimate stability boundary did not decrease enough to detect a drift in the estimate. Regardless, the high proportion of runs where no inattention was detected for *λ*_inattention_ = 0.25 does not present a strong limitation, as the estimation quality of uncorrected sequences suffers far less than for *λ*_inattention_ = 0.5.

The proposed method to correct for bias due to sustained inattention has several limitations. One major limitation is that sustained but short periods of inattention or intermittent inattention leading to less than four consecutive level increases will not be captured. The rule could be modified to *t*_PEST,3up_. However, preliminary simulations showed that the trade-off between an increased false positive rate and false negative rate using *t*_PEST,4up_ lead to an overall better estimation performance. On the other hand, combining the present method with a free lapse rate during the fitting process (Wichmann and Hill, 2001), allows addressing not only longer, but also shorter inattention periods or isolated lapses, while keeping the bias marginal. In the current implementation of the inattention correction method, if the PEST start level is chosen too low (i.e., below the threshold to be estimated), early level increases will occur, which could be interpreted as inattention. Although this has been addressed by not allowing *t*_PEST,4up_ ≤ 8, this may not suffice in every case. Yet, for PEST to be efficient, experimenters should have a rough idea of the order of magnitude of the threshold to be estimated, in order to choose an appropriate start level and start step. A more robust solution may involve modifying the stability boundary to be also a function of trial *t*. While the *t*_PEST,4up_ rule is specific to PEST, it could be adapted to other adaptive methods (e.g., staircase methods (Leek, 2001)). An advantage of the stability boundary is that it could be directly used in other (also non-adaptive) methods, as it does not depend on the psychophysical procedure, but only on the evolution of the threshold estimate 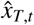.

It is possible that in some cases the estimation quality of parameters besides the threshold (e.g., slope or lapse rate) of the psychometric function may decrease when using the inattention correction method (e.g., in the example shown in **Figure 3**). However, PEST was only designed to converge towards the threshold and not to provide slope estimates (Taylor, 1971). There exist other sampling procedures specifically designed to estimate the slope or multiple parameters (e.g., (Levitt, 1971; Kontsevich and Tyler, 1999; King-Smith and Rose, 1997)).

Using the computer simulations, only a finite set of templates 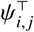 were tested and evaluated. However, they cover a range similar to the performance of healthy individuals in assessments of joint proprioception thresholds using various paradigms (Choi et al., 1995; Fry-Welch et al., 2003; Brewer et al., 2005; Tan et al., 2007; Lambercy et al., 2011; Rinderknecht et al., 2014; Elangovan et al., 2014; Cappello et al., 2015). Moreover, the variability in relative improvement across templates with different thresholds and slopes is one order of magnitude lower compared to the average relative improvement. Thus, the selected range of templates is representative enough to describe the performance of the inattention correction method.

In spite of the limitations, the proposed method goes beyond approaches identifying sequences of questionable validity such as the existing approaches by Hall (1983) and Leek et al. (1991). The results of the computer simulations demonstrated that, in contrast to other methods (Doll et al., 2015; Fründ et al., 2011), our method is capable of identifying specific inattention periods on a trial-by-trial basis, even within sequences of a small number of trials in a 2AFC experiment. Moreover, it allows correcting the data by removing these biased trials, leading to an improvement of estimates.

## 5 Behavioral data

As inattention is likely to be a strong confound in stroke patients (Tuhrim, 1993; Rinne et al., 2013) and will negatively affect assessment quality, the proposed method to correct for sustained inattention was tested on a set of previously recorded data from a proprioceptive assessment in this population. The experiment consisted of two sessions (test and retest, inter-session time span between 1 and 4 days). The joint angle difference threshold at the metacarpophalangeal joint of the index finger of both hands was assessed in both sessions using a one-degree-of-freedom robotic device (Rinderknecht et al., 2014).

As the “real” thresholds and whether or not the subjects suffered from sustained inattention are unknown (in contrast to the computer simulations), the threshold estimation errors could not be calculated. However, if a significant inattention period occurred during either the test or the retest, one can expect the discrepancy |Δ| between the two assessments to increase. We hypothesize that by applying the inattention correction, the difference between test and retest outcomes will be reduced in those cases where “inattention” was detected, as the biasing effect of a present inattention period would be reduced.

### 5.1 Methods

#### 5.1.1 Subjects

Data were collected from 20 stroke patients (> 2 weeks after their first clinical stroke, 10 right hemisphere stroke, 10 left hemisphere stroke, 12 male, 8 female, age mean ± SD: 66.8 ± 8.3 years). Besides the inability to detect large passive finger movements, exclusion criteria comprised severe hand edema, high muscle tone, or pain, as it would hinder the correct use of the assessment device. Further exclusion criteria were severe cognitive impairment, aphasia and neglect. Eighteen subjects were right handed and two were ambidextrous, according to the Edinburgh Handedness Inventory (Oldfield, 1971). In accordance with the Declaration of Helsinki, all subject gave written informed consent prior to their participation. The study was approved by the institutional ethics committee of the University of Konstanz.

#### 5.1.2 Procedure

A two-interval 2AFC paradigm was used in combination with PEST with a logarithmic domain adaptation (Rinderknecht et al., 2014). In each trial, two consecutive passive flexion movements were presented by the robotic device, both starting from a neutral position (i.e., 0°, hand aligned with forearm). Each flexion movement lasted 1 s, independent of the movement amplitude. The finger remained at the tested angle for 1.5 s, before moving back to neutral position at the same speed. After the presentation of the two different angles, the subject was asked to indicate which of the two angular movements was larger by providing their response directly with their other hand on a touchscreen. PEST defined the angular difference (referred to as level) between the two intervals. Each pair of presented angles was symmetrically arranged around 20°, and the order of larger versus smaller angle was randomized. By limiting the movements to flexion, the stimulus range was constrained to [0°, 40°]. The same PEST parameters and termination conditions as in the computer simulations were used (start level of 5.5°, start step of 2°, *P*_*t*_ of 75%, *W* of 1). Termination conditions were (i) a maximum number of 60 trials, (ii) 20 consecutive trials at the same level, or (iii) a minimum step of 0.1°. The tested limb was occluded from vision during the entire experiment. In each of the two sessions, proprioception of both hands was assessed in randomized order.

#### 5.1.3 Data analysis

For each test and retest pair the absolute difference |Δ| between the two difference threshold estimates was calculated for both conditions, without and with inattention correction. These differences |Δ| were pooled across subjects and assessed hands. The level of impairment, as well as whether the contra- or ipsilesional hand was tested, was irrelevant for the purpose of evaluating the inattention correction method. Furthermore, no inferences were made about the performance of proprioceptive discriminability of the stroke patients, and assessing and pooling both hands allowed a larger dataset with subjects being likely to suffer from inattention. All analyses were conducted in MATLAB R2014a.

### 5.2 Results

Inattention was detected in 11 out of 80 assessment sequences. In those 11 sequences, 18 ± 8 trials were rejected on average, and the exclusion period started on average at trial 33 ± 9 (onset *t*_[exclude_). In 6 of those sequences the inattention period ended before the end of the assessment, whereas in the other 5 sequences the end of the assessment was reached before. Thus, in the latter cases only data of trials up to the inattention onset were used for the threshold estimate when applying the inattention correction method. **Figure 8** illustrates the absolute differences |Δ| between test and retest for both conditions without and with inattention correction plotted against each other (left) and their distributions (right). In 8 sequence pairs |Δ| was decreased (7/11 with an absolute change ≥ 0.1°, corresponding to the minimum step of the PEST algorithm), while in 3 sequence pairs |Δ| increased (2/11 with an absolute change ≥ 0.1°).

**Figure 8.**
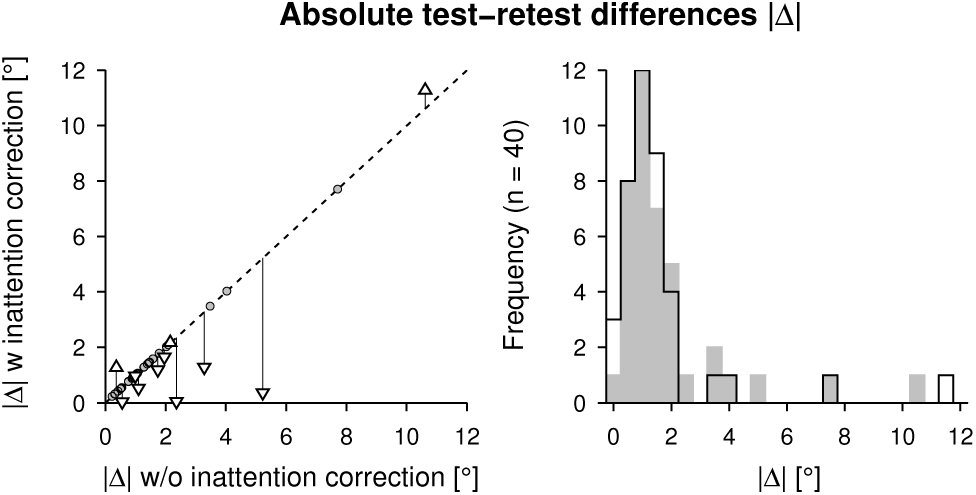
Behavioral test-retest data from the proprioceptive assessment conducted for both hands in 20 stroke patients. (Left) Absolute test-retest differences |Δ| for no inattention correction vs. when the algorithm was applied to detect and correct for sustained inattention. Test-retest differences from sequences which were not modified by the inattention correction method are indicated by circles. Where the algorithm had an effect on the threshold estimates, the |Δ| is indicated by triangles pointing downwards (decreased |Δ| with inattention correction) or upwards (increased |Δ| with inattention correction) connected by black vertical lines to the dashed diagonal identity line to better emphasize the change. In 8 of 11 inattention cases the test-retest discrepancy could be reduced by applying the inattention correction method. **(Right)** Comparison of |Δ| -distributions (bin resolution of 0.5°) without (gray fill) and with (black outline) inattention correction. The overall decrease of variability when applying the inattention correction method is visible as the shift of the distribution towards |Δ| = 0°.

### 5.3 Discussion

The aim of applying the inattention correction algorithm on a real dataset from a behavioral experiment was to explore whether the proposed method would detect “inattention” in non-simulated data and whether it could decrease the testretest discrepancy |Δ| in those cases, by excluding potentially biased data. Indeed, the algorithm detected “inattention” in 14% of the assessments. Correcting the data by removing trials during inattention periods provided more similar testretest assessment results in 73% of the corrected assessment test-retest pairs. In the other few cases, the difference between test and retest was only marginally augmented (on average by 0.53°).

It is noteworthy that the two largest discrepancies were not reduced with the inattention correction method. Detailed analysis of the PEST sequences revealed that in one case the PEST sequence increased by 3 steps right from the beginning before the PEST level decreased at trial 22. Due to the early increase, the PEST algorithm could not fully converge (defined by minimum step termination criterion) within the remaining trials of the assessment. What looked like suboptimally chosen start parameters of the sampling procedure, may have resulted from initial inattention or unfamiliarity with the task (e.g., the task may not have been fully understood), affecting performance, as the retest of the same hand showed immediate convergent behavior towards a threshold lower than the start level. Hence, it may be advisable to either increase the start level even more or conduct a few familiarization trials before the actual assessment.

In the other case of large |Δ|, “inattention” was detected in the first of the two test-retest assessments and the difference threshold was reduced by about 15% by the algorithm. In the retest, no “inattention” was detected, even though experienced experimenters would suggest an inattention period when visually inspecting the PEST sequence. Indeed, there are 6 consecutive PEST level increases. However, it was not captured by the inattention detection algorithm due to the stability boundary. When manually removing the visually identified “inattention” period, |Δ| decreased from around 10.6^°^ to 1.7°. Further tuning or a decrease of the stability boundary width would allow detecting such a case, but to the detriment of false negative detections.

Coincidently, the average detected inattention onset and exclusion period length in this dataset with stroke patients were very similar to the values used for the inattention models in the computer simulations. Consequently, this might suggest that the way inattention was modeled in the computer simulations (onset distribution and inattention period) corresponds to a realistic scenario, despite being a simplified model (e.g., constant inattention level during the entire inattention period). Furthermore, considering that the task in the present proprioceptive assessments is highly repetitive and that stroke patients may suffer from increased fatigue (Staub and Bogousslavsky, 2001; de Groot et al., 2003; Duncan et al., 2012) and inattentiveness (Tuhrim, 1993; Rinne et al., 2013), it seems reasonable that some subjects start losing attention to the task after 5–10 minutes (around trial 30), also considering literature pointing at attention spans of 7–20 minutes in students following lectures (Bligh, 1998; Petty, 2004; David and Dukette, 2009; Dent and Harden, 2013). Contributing factors to inattention during the proprioceptive assessment may also be (i) increased familiarity with the task and (ii) increased task difficulty (as the adaptive sampling procedure converges towards the threshold) leaving the subject with the potentially uncomfortable impression that he/she is guessing and thus uncertain about the correctness of the answer (Treutwein, 1995; Kaernbach, 2001).

Considering restrictions for administration time of clinical assessments in stroke patients (Gresham et al., 1996), or in clinical populations in general, only assessments with a small number of trials can be integrated into a clinical setting, for which reason the current proprioceptive assessment was limited to 60 trials. Based on the behavioral results with stroke patients, the average end of the inattention period was around trial 50 ± 11, which is close to the end of the assessment. In the computer simulations it could be shown that, when the inattention correction is not applied, the estimation quality is poorest if the assessment is stopped right after the end of the inattention period, as the ratio of biased versus unbiased data is the largest. Thus, using the proposed method to address inattention confounds might be even more essential in such scenarios where the assessments have to be short.

## 6 Conclusions

Loss of attention presents challenges in proprioceptive threshold testing which have not been sufficiently addressed. For instance, inattention can affect perception during sensory assessments and bias psychophysical threshold estimates. We proposed a method capable of efficiently identifying periods of sustained inattention based on the recorded stimulus-response pairs of a PEST sequence together with the evolution of the estimated threshold. This algorithm allows excluding specific trials with biased data due to inattention and improving the threshold estimates, as evaluated in computer simulations and tested on a set of behavioral data, which has important implications for the psychometric properties of an assessment, such as reliability and validity. The method can also be used for sequences with relative small numbers of trials and in experiments using a 2AFC paradigm. Furthermore, the inattention correction method complements existing methods addressing lapses and goodness of fit (e.g., freeing up the lapse rate of the psychometric function fit (Wichmann and Hill, 2001)), such that not only isolated lapses but also longer inattention periods can be addressed to improve the quality of the outcome measure. The proposed method can be adjusted to other adaptive procedures (e.g., staircase), and can be used both offline to correct previously recorded experimental data *post-hoc*, as well as online (with some limitations) to generate an alert of inattention, and pause or stop the experiment to avoid further collection of biased data. When resuming the experiment, the sampling procedure could continue at the stimulus level at which the inattention onset was detected. This may allow shortening experimental procedures, which could have a significant impact on the improvement of assessment practice, especially in studies with patient populations potentially suffering from attention deficits and where experimental time is costly, while providing more reliable estimates. In addition, it could be shown that the benefit of correcting for biases is even higher when assessments are short, as in these cases a sustained inattention period has a larger detrimental effect on the estimation quality. While the statistics based on the results of the computer simulations have shown that there is an overall improvement of estimation quality at the group level when applying inattention correction, it can also positively impact diagnosis, prognosis, or treatment planning for the individual patient if inattention confounds can be removed from the assessment’s outcome measures.

## Author contributions

MR, RR, WP, OL and RG contributed to the conception of this work. MR and RR developed the methodology. MR implemented the experiments and the computer simulations, performed the analysis, interpreted the results, and drafted the manuscript. MR, RR, WP, OL and RG revised the manuscript and approved the final version.

## Acknowledgments

The authors would like to thank J. Liepert and V. Raible from the Kliniken Schmieder Allensbach for their collaboration and acquisition of the behavioral data, as well as K. Leuenberger for fruitful discussions regarding the development of the algorithm. This research was supported by the ETH Zurich Foundation in collaboration with Hocoma AG, by the Janggen-Pöhn Foundation, and the Schmieder Foundation for Science and Research.

